# High diversity in Delta variant across countries revealed via genome-wide analysis of SARS-CoV-2 beyond the Spike protein

**DOI:** 10.1101/2021.09.01.458647

**Authors:** Rohit Suratekar, Pritha Ghosh, Michiel J.M. Niesen, Gregory Donadio, Praveen Anand, Venky Soundararajan, A.J. Venkatakrishnan

**Author notes:** Correspondence to: AJ Venkatakrishnan and Venky Soundararajan. Equal contribution (Rohit Suratekar, Pritha Ghosh).

## Abstract

The highly contagious Delta variant of SARS-CoV-2 has emerged as the new dominant global strain, and reports of reduced effectiveness of COVID-19 vaccines against the Delta variant are highly concerning. While there has been extensive focus on understanding the amino acid mutations in the Delta variant ‘s Spike protein, the mutational landscape of the rest of the SARS-CoV-2 proteome (25 proteins) remains poorly understood. To this end, we performed a systematic analysis of mutations in all the SARS-CoV-2 proteins from nearly 2 million SARS-CoV-2 genomes from 176 countries/territories. Six highly-prevalent missense mutations in the viral life cycle-associated Membrane (I82T), Nucleocapsid (R203M, D377Y), NS3 (S26L), and NS7a (V82A, T120I) proteins are almost exclusive to the Delta variant compared to other variants of concern (mean prevalence across genomes: Delta = 99.74%, Alpha = 0.06%, Beta = 0.09%, Gamma = 0.22%). Furthermore, we find that the Delta variant harbors a more diverse repertoire of mutations across countries compared to the previously dominant Alpha variant (cosine similarity: mean_Alpha_ = 0.94, S.D._Alpha_ = 0.05; mean_Delta_ = 0.86, S.D._Delta_ = 0.1; Cohen ‘s d_Alpha-Delta_ = 1.17, p-value < 0.001). Overall, our study underscores the high diversity of the Delta variant between countries and identifies a list of targetable amino acid mutations in the Delta variant ‘s proteome for probing the mechanistic basis of pathogenic features such as high viral loads, high transmissibility, and reduced susceptibility against neutralization by vaccines.

## Introduction

The ongoing COVID-19 pandemic has infected over 210 million people and killed nearly 4.5 million people worldwide as of August 2021^1^. Throughout the pandemic, the SARS-CoV-2 virus has acquired novel mutations, and the US government SARS-CoV-2 Interagency Group (SIG) has classified the mutant strains as Variant of Concern (VOC), Variant of Interest (VOI), and Variant of High Consequence (VOHC)^2^. The variants of concern (Alpha, Beta, Gamma, and Delta) are more transmissible, cause more severe disease, and/or reduce neutralization by vaccines and monoclonal antibodies^2,3^. The Delta variant (PANGO lineage B.1.617.2), first isolated from India in October 2020^3^, has emerged as the dominant global variant alongside the Alpha variant (PANGO lineage B.1.1.7), with genome sequences deposited from 104 and 150 countries, respectively, in the GISAID database^4^ and has worsened the public health emergency^5,6^.

Recent studies are reporting nearly 1000-fold higher viral loads in infections associated with the Delta variant^7^ and reduced neutralization of this variant by vaccines^8–12^. The NCBI database lists 26 proteins (structural, non-structural, and accessory proteins) in the SARS-CoV-2 proteome^13^ totaling 9757 amino acids. These include four structural proteins (Spike, Envelope, Membrane, and Nucleocapsid), 16 non-structural proteins (NSP1–NSP16), and six accessory proteins (NS3, NS6, NS7a, NS7b, NS8, and ORF10). As of August 2021, the CDC identifies 11 amino acid mutations in the Spike protein of the Delta variant^2^, and the functional role of the SARS-CoV-2 Spike protein mutations has been well studied^14–16^. However, the mutational landscape of the rest of the Delta variant ‘s proteome remains poorly understood. Concerted global genomic data sharing efforts through the GISAID database^4^ have led to the availability of nearly 2 million SARS-CoV-2 genomes from over 175 countries/territories, thereby providing a timely opportunity to analyze the mutational landscape of SARS-CoV-2 variants across all the 26 proteins.

Here, we perform a systematic analysis of amino acid mutations across the SARS-CoV-2 proteome (26 proteins) for the variants of concern and identify that the Delta variant harbors the highest mutational load in this proteome. Interestingly, the Delta variant ‘s proteome is also highly diverse across different countries compared to the Alpha variant. Our observations suggest the need to account for country-specific mutational profiles for comprehensively understanding the biological attributes of the Delta variant such as increased viral loads and transmissibility, and reduced susceptibility against neutralization by vaccines.

## Results

### Delta variant has highly prevalent mutations in the viral life cycle-associated Membrane, Nucleocapsid, NS3, and NS7a proteins

Currently, only the Spike protein mutations are being used in literature to define the SARS-CoV-2 variants of concern and interest^2,3^. However, the analysis of 1.99 million genome sequences of SARS-CoV-2 from 176 countries/territories in the GISAID database^4^ revealed mutations in 52.3% of the 9757 amino-acid-long SARS-CoV-2 proteome. In all, there are 8157 unique mutations in 5107 amino acids spanning 24 of the 26 SARS-CoV-2 proteins (**Figure S1**). The Spike protein contributes to only 6.3% (1055 unique mutations in 617 amino acids) of the mutated SARS-CoV-2 proteome, which emphasizes the need to study the mutational profile across all the proteins of SARS-CoV-2.

Of the 1.99 million SARS-CoV-2 genomes analyzed here, there are 198,460 genomes corresponding to the Delta variant from 104 countries. We identified seven highly prevalent mutations in the following proteins of the Delta variant: Membrane (I82T: 99.9%), Nucleocapsid (R203M: 99.9%, D377Y: 99.6%), NSP12 (P323L: 99.9%), NS3 (S26L: 99.9%), and NS7a(V82A: 99.4%, T120I: 99.7%). Strikingly, all these mutations except P323L in NSP12 are nearly exclusive to the Delta variant compared to other variants of concern (mean prevalence_Delta_ = 99.74%, mean prevalence_otherVariantsofConcern_ = 0.12%) (**Figure S2, Table S1**). Within the Spike protein, there are four such mutations (T19R, L452R, T478K, P681R) as well (mean prevalence_Delta_ = 99.86%, mean prevalence_otherVariantsofConcern_ = 0.04%). In total, there are ten mutations across the proteome that are characteristic of the Delta variant. This list of highly prevalent mutations can serve as candidates for probing the mechanistic basis of the Delta variant ‘s pathogenic features such as high viral loads, increased transmissibility, and reduced susceptibility against neutralization by vaccines. Indeed, the I82T mutation of the SARS-CoV-2 Membrane protein of the Delta variant has been proposed previously to increase biological fitness with altered glucose uptake during viral replication^17^. The known functional implications of Delta variant mutations include antibody escape^18–21^, high viral load^22^, increased transmissibility^7,23^, and infectivity^24^ (**Table 1**).

**Table 1:** Functional implications of mutations in SARS-CoV-2 Delta variant. Mutations in the SARS-CoV-2 Delta variant with known functional implications.

| Mutation | Functional domain/region | Is solvent-accessible? | Functional implications |
| --- | --- | --- | --- |
| Spike_E156G | N-terminal domain | Yes | Antibody escape<sup>18,19</sup> |
| Spike_ΔF157 | N-terminal domain | Yes | Antibody escape<sup>18,19</sup> |
| Spike_ΔR158 | N-terminal domain | Yes | Antibody escape<sup>18,19</sup> |
| Spike_L452R | Receptor-binding domain | Yes | Antibody escape<sup>20,21</sup> |
| Spike_T478K | Receptor-binding domain | Yes | Antibody escape<sup>20,21</sup> |
| Spike_D614G | - | Yes | Increases spike density and infectivity of virion<sup>24</sup>, and viral replication<sup>22</sup> |
| Spike_P681R | - | Yes | Increased transmissibility<sup>23,37</sup> |
| M_I82T | Membrane-spanning helix (TM3)<sup>17</sup> | Yes | More biologically fit, with altered glucose uptake during viral replication<sup>17</sup> |
| NSP12_P323L | - | Yes | Increased transmissibility<sup>38</sup> |

### Delta variant is variable across countries and has country-specific core mutations

While the Alpha variant spread widely during the pre-vaccination phase of the pandemic^3,25^, the Delta variant emerged as a global strain during the vaccination period. Given that the extent of vaccination coverage is highly variable across countries^26^, the selection pressure against the Delta variant is also likely to vary. To understand mutational profiles of SARS-CoV-2 variants of concern across countries, we generate ‘mutational prevalence vectors ‘ for each country of occurrence and calculate their pairwise cosine similarities (**Figure 1A**, *Methods*). The cosine similarity distributions for the Alpha and Delta variants are significantly different (Jensen-Shannon divergence = 0.21, 95% confidence Interval: [0.17, 0.24], p-value < 0.001). The mean and standard deviation (S.D.) of pairwise cosine similarity values for the globally dominant Alpha and Delta variants (mean_Alpha_ = 0.94, S.D._Alpha_ = 0.05; mean_Delta_ = 0.86, S.D._Delta_ = 0.1) show a significantly higher diversity in the Delta variant as compared to Alpha (Cohen ‘s d = 1.17, 95% confidence Interval: [1.02, 1.28], p-value < 0.001) (**Figure 1B, Figure S3**).

**Figure 1:**
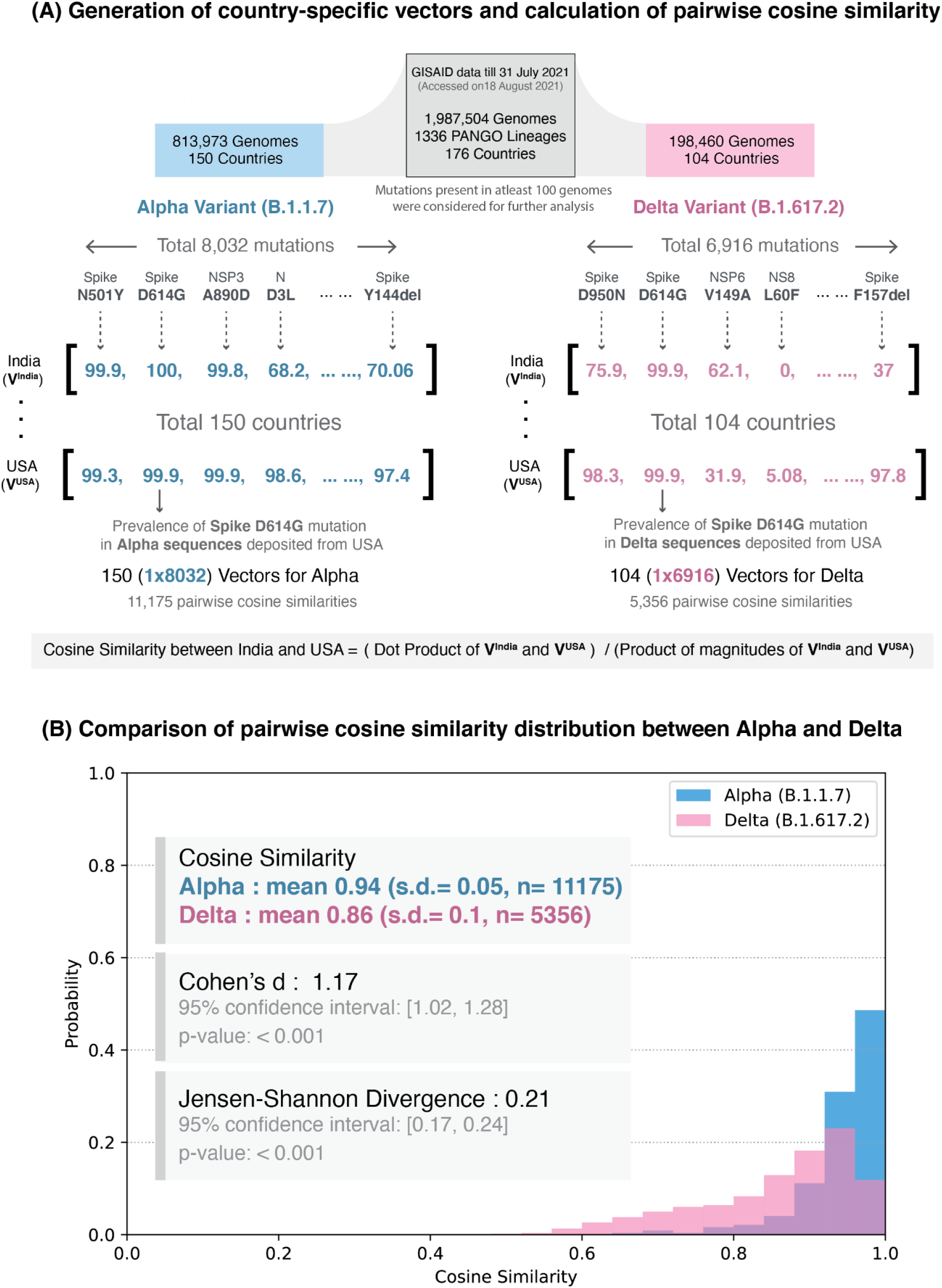
Schematic overview of the study. **(A)** Generation of country-specific mutation prevalence vectors and calculation of pairwise cosine similarity. The study dataset, updated as of 31 July 2021, with nearly 2 million sequences were retrieved from GISAID. For a variant of concern, mutational prevalence vectors were calculated for each country of their occurrence. For example, the Delta variant has been reported in 104 countries worldwide and harbors 6916 unique mutations. Thus we generate 104 mutational prevalence vectors with (1 × 6916) dimensions and calculate the pairwise cosine similarities for ^104^C_2_ (5356) combinations. **(B)** Comparison of probability distributions of pairwise cosine similarity values for the Alpha and Delta variants. The cosine similarity distributions for the Alpha and Delta variants are significantly different (Jensen-Shannon divergence = 0.21, 95% confidence Interval: [0.17, 0.24], p-value < 0.001). The mean and standard deviation (s.d.) of pairwise cosine similarity values for the globally dominant Alpha and Delta variants show a significantly higher diversity in the Delta variant as compared to Alpha (Cohen ‘s d = 1.17, 95% confidence Interval: [1.02, 1.28], p-value < 0.001).

To determine mutations that can contribute to country-specific differences in the Delta variant, we identified the highly-prevalent mutations at the country level (“country-specific core mutations”) (**Figure 2A**; *Methods*). As an example, here we compare the country-specific core mutations in the United States (Delta_UnitedStates_) and in India (Delta_India_). Delta_UnitedStates_ has 29 country-specific core mutations compared with 19 country-specific core mutations in Delta_India_ (**Figure 2B**). Of these, 16 mutations are common, spanning structural proteins (Spike, Nucleocapsid, and Membrane), non-structural proteins (NSP3, NSP4, NSP6, NSP12, and NSP13), and accessory proteins (NS3, and NS7a). There are three mutations in three proteins that are highly prevalent in Delta_India_ but not in Delta_Unitedstates_. In contrast, there are 13 mutations spanning six proteins that are highly prevalent in Delta_UnitedStates_ but not Delta_India_, including in the exoribonuclease NSP14, which is critical for the viral replication machinery^27^ and can inhibit the host translational machinery^28^. The presence of country-specific differences in the Delta variants motivate the need to understand whether these genome-level differences manifest differences in the disease phenotypes and vaccine effectiveness.

**Figure 2:**
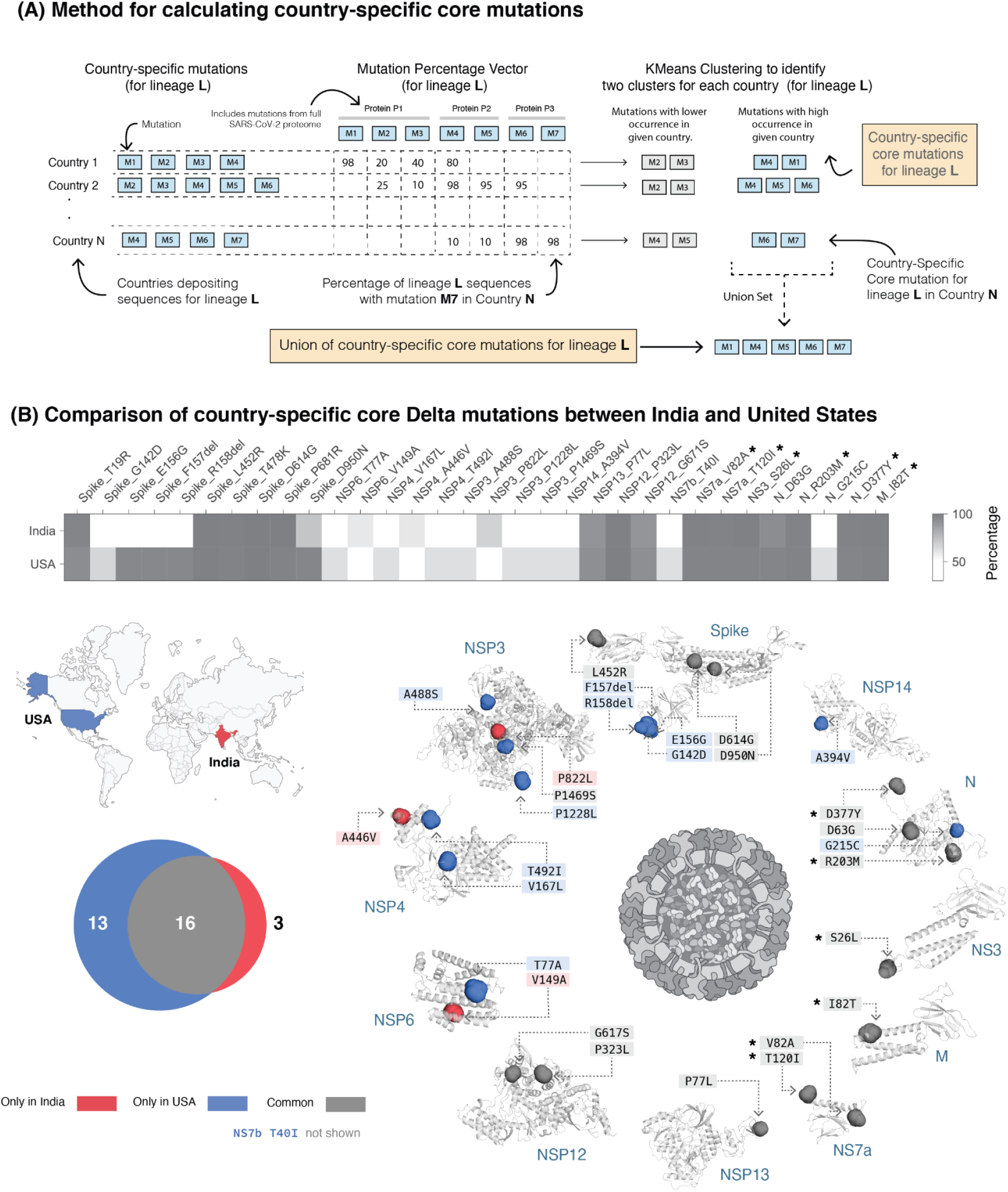
Identification of country-specific core mutations. **(A)** Schematic overview of the method for defining country-specific core mutations for a lineage. See *Methods* for further details. **(B)** Comparison of prevalence of country-specific core mutations in the Delta variant in India and the United States. Mutations marked with an asterisk are highly prevalent in all countries of occurrence of Delta variant (mean prevalence = 99.28%) but are nearly absent(mean prevalence = 2.62%) in the other variants of concern. The Venn diagram shows the intersection and union set of country-specific core mutations among the two countries. The mutations are highlighted (in the same color scheme as the Venn diagram) on the structure of the Spike protein and the structural models of the other SARS-CoV-2 proteins (see *Methods*). Residues corresponding to Spike protein mutations T19R, T478K, and P681R are missing from the structure of the Spike protein and hence not shown here. The 43-amino acid long NS7b protein has no structure/model available and hence is not represented here.

## Discussions

COVID-19 is the first pandemic of the post-genomic era^29^ that has been under intense genomic surveillance through concerted global viral sequencing efforts. This has led to the identification and tracking of emerging variants of concern, such as the highly transmissible Alpha variant and Delta variant. Through analysis of nearly 2 million genomes from 176 countries/territories, we have identified that there are mutations beyond the Spike protein that are characteristic of the Delta variant and that the Delta variant is more variable across countries than other variants of concern.

Our study has identified ten highly prevalent mutations characteristic of the Delta variant across five proteins, which can serve as therapeutic targets and as candidates for probing the mechanistic basis of the Delta variant ‘s pathogenic features such as high viral loads, increased transmissibility, and reduced susceptibility against neutralization by vaccines. The country-specific differences in the Delta variant ‘s mutational profile identified in this study can also be used to guide the design of vaccines/boosters that can comprehensively combat COVID-19. Our study also motivates that the diversity at the proteome level should be considered in designating the variants of concern and interest. Future studies are warranted to comprehensively examine the combinations of factors such as vaccination rates, geographical proximity (**Figure S4**), and air connectivity (**Figure S5**) to dissect the difference in the epidemiology of Delta variants across countries.

This study has a few limitations. Since this study is based on publicly available data from the GISAID database, it may carry biases associated with sequencing disparities across countries and reporting delays. Though there is extensive genomic surveillance, there is a lack of clinical annotation of the genomes, limiting our ability to assess the clinical impact of the country-specific differences in the variants. The GISAID database does not record mutations in the recently discovered ORFs in the SARS-CoV-2 genome such as ORF10, ORF9b, and ORF9c. The assignment of the mutations in these ORFs may reveal further differences between SARS-CoV-2 variants.

Though mass vaccination efforts are underway around the world, there are huge differences in the population immunity of countries due to the differences in the vaccines approved regionally and the extent of vaccination coverage in populations. These differences contribute to the risk of emergence of new SARS-CoV-2 variants and continued genome-surveillance is imperative for developing comprehensive global and country-specific preventive and therapeutic measures to end the ongoing pandemic.

## Methods

### SARS-CoV-2 genome sequences

We retrieved 1,987,504 SARS-CoV-2 high coverage complete genome sequences from human hosts in 176 countries/territories spanning 1336 PANGO lineages on 18 August 2021 from GISAID^4^ for December 2019 - July 2021, of which 816 sequences do not harbor any mutations. We removed sequences from other hosts and those with incomplete dates (YYYY-MM or YYYY) from further analyses. 1,986,688 sequences harbor a total of 89,875 unique mutations. However, to account for errors arising from sequencing, we only consider 8157 unique mutations in 24 proteins that are present in 100 or more sequences for all our further analyses. We did not identify any mutations in NSP11 (for which no mutations are present in 100 or more sequences) and ORF10 (for which no information on mutations are available in GISAID data), and hence are not considered in further analyses.

Though 99.15% of all SARS-CoV-2 genome sequences possess one or more mutations in the Spike protein, 98.91% and 95.2% of sequences also bear mutations in the crucial NSP12 (RNA-dependent RNA polymerase, RdRp) and Nucleocapsid proteins, respectively.

We retrieved the list of proteins in the SARS-CoV-2 proteome from NCBI^30^ on 2 August 2021. The structure of the Spike protein was retrieved from PDB (code: 6VSB) and that of the structural models of the other SARS-CoV-2 proteins from https://zhanglab.ccmb.med.umich.edu/COVID-19/ (on 11 June 2021).

### Cosine similarity across countries

To calculate the cosine similarity of a lineage *L* among countries, we generated a prevalence vector of constituent mutations for each country of occurrence of the lineage *L*. For a pair of countries, the cosine similarity of the lineage *L* was calculated for their mutation vectors (A, B) (**Equation 1, Figure 1A**).

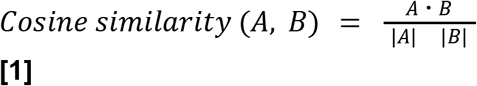

The mean and standard deviation (S.D.) of pairwise cosine similarity values for variants of concern (mean_Alpha_ = 0.94, S.D._Alpha_ = 0.05; mean_Beta_ = 0.89, S.D._Beta_ = 0.06; mean_Gamma_ = 0.95, S.D._Gamma_ = 0.03; mean_Delta_ = 0.86, S.D._Delta_ = 0.1) show a higher diversity of the Delta variant across countries. To check the effect size, Cohen ‘s d was calculated (**Equation 2**).

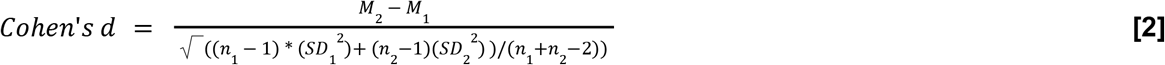

where, *M*: mean, *n*: sample size, *SD*: standard deviation

Probability distributions of pairwise cosine similarities were calculated by binning frequencies (bins = 25), and their Jensen-Shannon divergence (with base 2) was calculated using the *jensenshannon* function available in SciPy [v1.7.0]^31^. p-value was calculated using bootstrapping with 1000 iterations.

To identify countries with similar mutational profiles, we clustered the pairwise cosine similarity matrix with Ward ‘s variance minimization algorithm^32^ available in SciPy [v1.7.0].

### Bootstrapping of cosine similarities

For each country, we resampled (with replacement) all the sequences deposited in the GISAID database and generated a cosine similarity distribution for Alpha and Delta variants (**Figure S6**). For calculating 95% confidence interval, we calculated Jensen-Shannon divergence (JSD) and Cohen ‘s d for each bootstrap iteration. To get a null distribution for JSD and Cohen ‘s d, we calculated these metrics from the Alpha and Delta cosine similarity distribution generated in each bootstrap iteration (n = 1000). The p-values were calculated based on the distribution of all bootstrapped values and original JSD / Cohen ‘s d values.

### Cosine similarity for airline connectivity

Air traffic data was accessed on 13 June 2021 from The OpenSky Network 2020^33,34^. Only international flights were considered in this analysis. A matrix of the number of flights across all countries of the world was generated for the period of February 2021 - June 2021. For country *A*, a vector of the number of outgoing flights to all the other countries normalized with respect to the total number of outgoing flights from country *A* was generated. Similarly, for country *B*, a vector of the number of incoming flights from all the other countries normalized with respect to the total number of incoming flights to country *B* was generated. Cosine similarity for airline connectivity for this pair of countries was calculated as in **Equation 1**.

### Country-specific core mutations

Genome sequences of Alpha, Beta, Gamma, and Delta variants in GISAID data are available from 150, 95, 61, and 104 countries, respectively. For country *C*, we calculated the prevalence of a mutation *M* as in **Equation 3**.

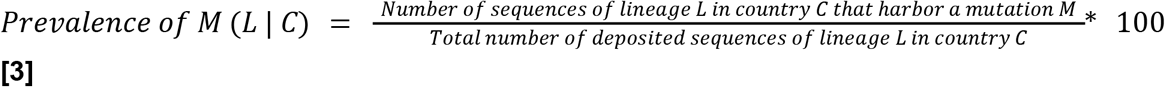

The prevalence of all mutations identified in lineage *L* in country *C* was calculated and further clustered using K-means clustering algorithm^35^ (in scikit-learn^36^) for unbiased identification of the highly prevalent set (core) of mutations for lineage *L* in country *C*. Based on K-means clustering sensitivity analysis, we partitioned the observations into two clusters for K-means clustering with initial cluster centroids at 0% and 100% (**Figure S7**). All mutations with labels corresponding to the higher centroid are called the core mutations of lineage *L* in country *C* (“country-specific core mutations”). A union set of country-specific core mutations from all countries in which lineage *L* is present were also determined. We observed that the Delta variant ‘s union set of country-specific core mutations are distinct and higher from those in the other variants of concern (**Figure S8, Table S2**).

The characteristic Spike protein mutations defined by the CDC^2^ (as of 2 August 2021) overlap with those identified in our analysis (**Figure S9**), thus validating our method of identifying mutations in the SARS-CoV-2 proteome.

## Acknowledgements

The authors thank Murali Aravamudan, Arjun Puranik, Sutirtha Chakraborty, Gajinder Pal Singh and Shahir Asfahan for feedback on this manuscript.

## Author Contributions

VS and AJV conceived the study. PG, RS, MJMN and AJV designed the study, reviewed the findings and wrote the manuscript. RS, PG, MJMN, GD, PA, AJV and VS contributed methods, data, analysis, or software. All authors revised the manuscript.

## Supplementary Information

**Figure S1:**
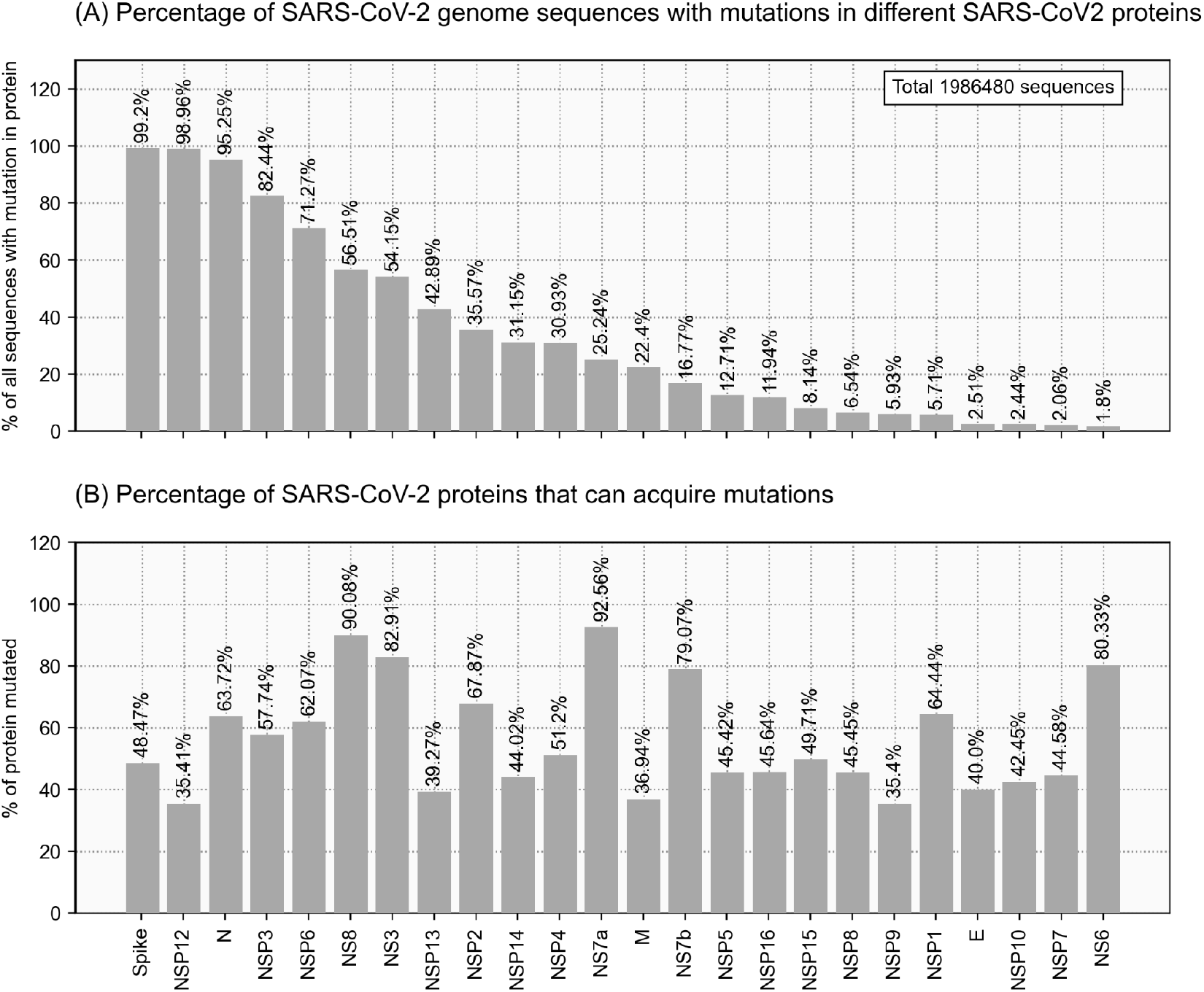
The proteome-wide mutational profile of SARS-CoV-2. **(A)** Percentage of SARS-CoV-2 genome sequences with mutations in different SARS-CoV-2 proteins. Spike, NSP12, and Nucleocapsid are the most frequently mutated proteins. **(B)** Percentage of SARS-CoV-2 proteins that can acquire mutations. The NS7a and NS8 are hypervariable proteins that mutate 92.56% and 90.08% of their sequence lengths.

**Figure S2:**
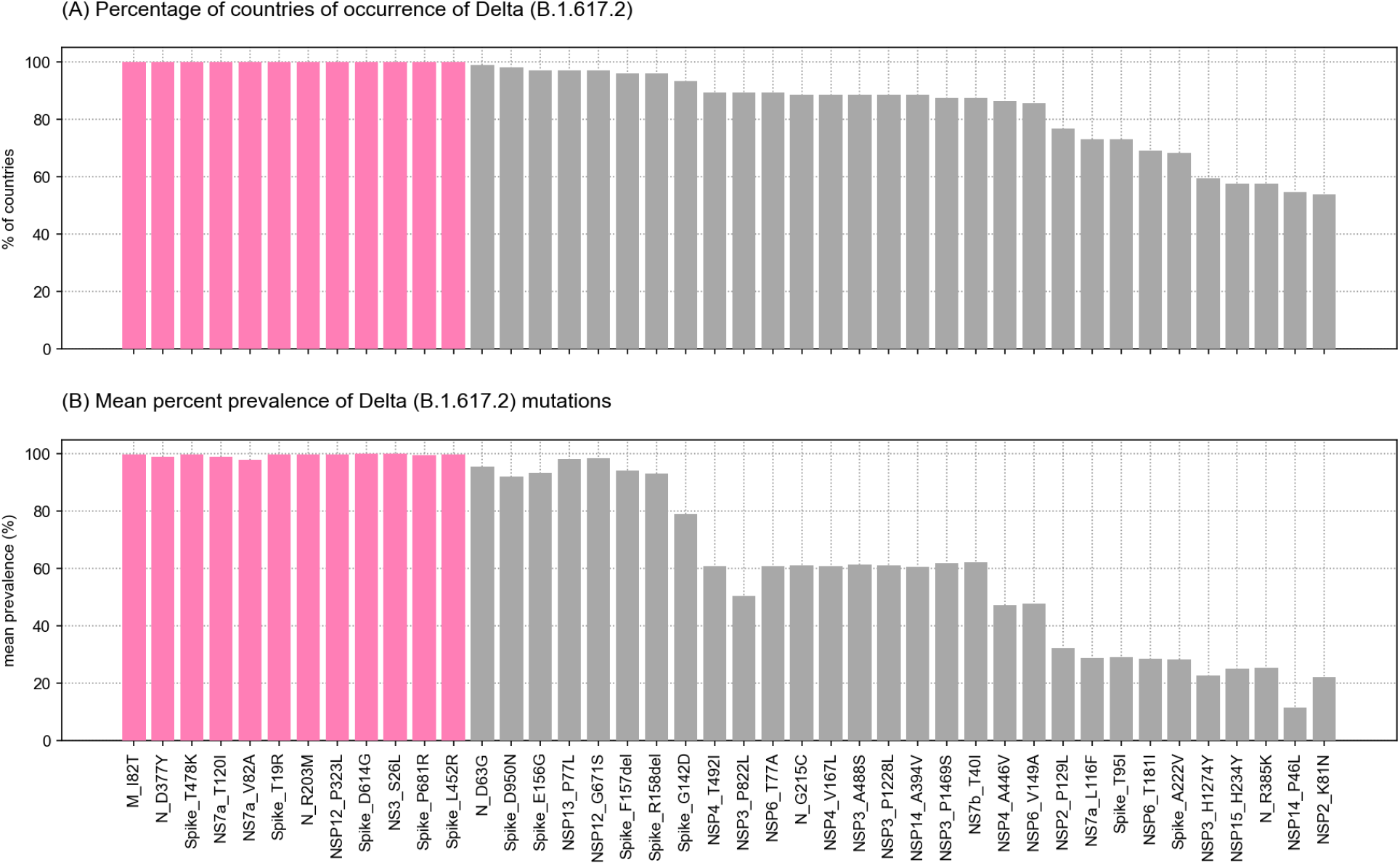
Prevalence of mutations in the Delta variant. **(A)** Percentage of countries of occurrence of the Delta variant. Only mutations present in 50% or more countries are shown here for representative purposes. 12 mutations (highlighted in magenta) are reported from all 104 countries where the Delta variant occurs. **(B)** Mean percent prevalence of Delta mutations. The 12 mutations highlighted in magenta are highly prevalent (mean prevalence range: 97.99 - 99.94%) in all countries of occurrence of the Delta variant.

**Figure S3:**
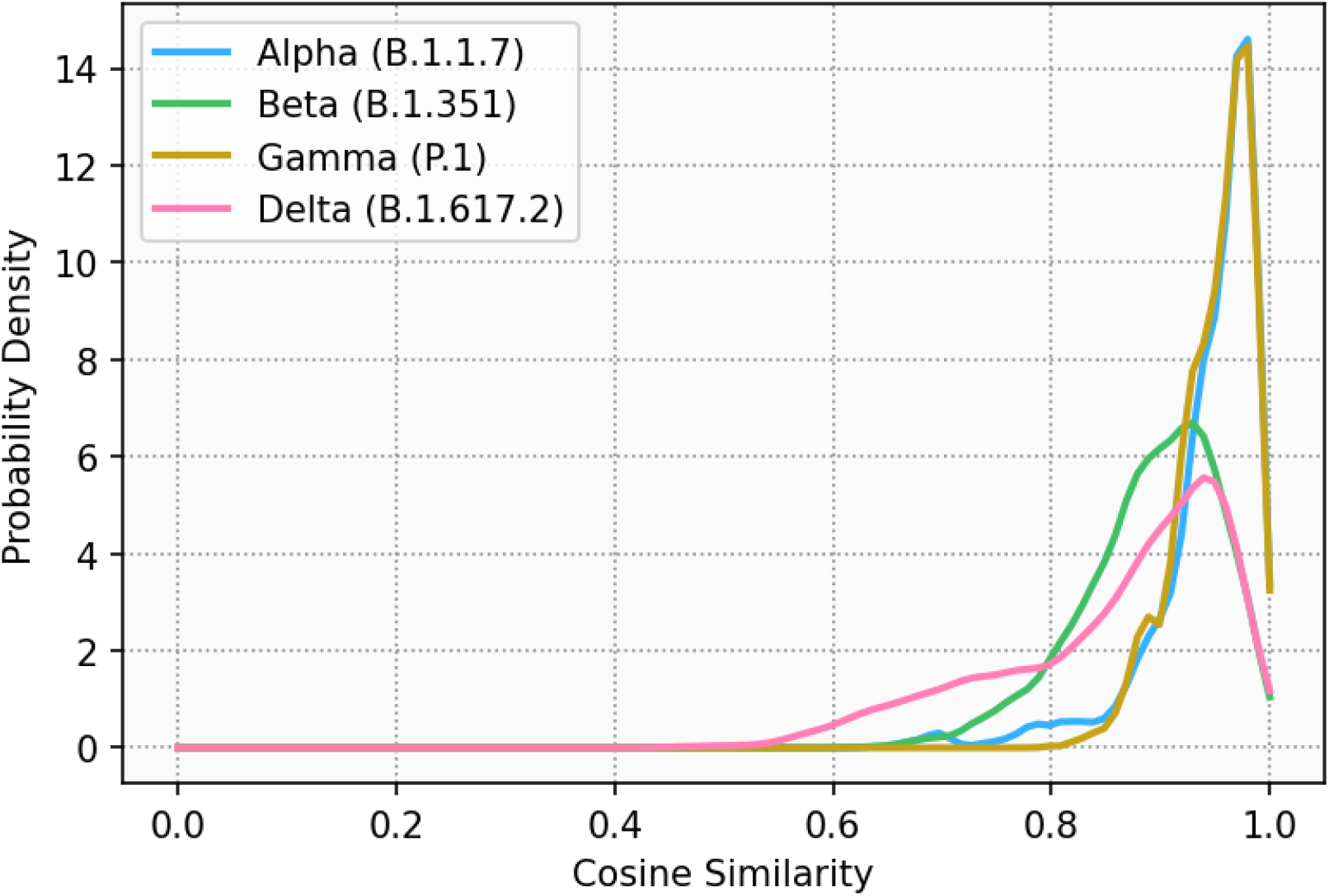
Probability density of pairwise cosine similarity across countries. The plot suggests a higher diversity of pairwise cosine similarity values, and thus a higher diversity of the Delta variant across countries.

**Figure S4:**
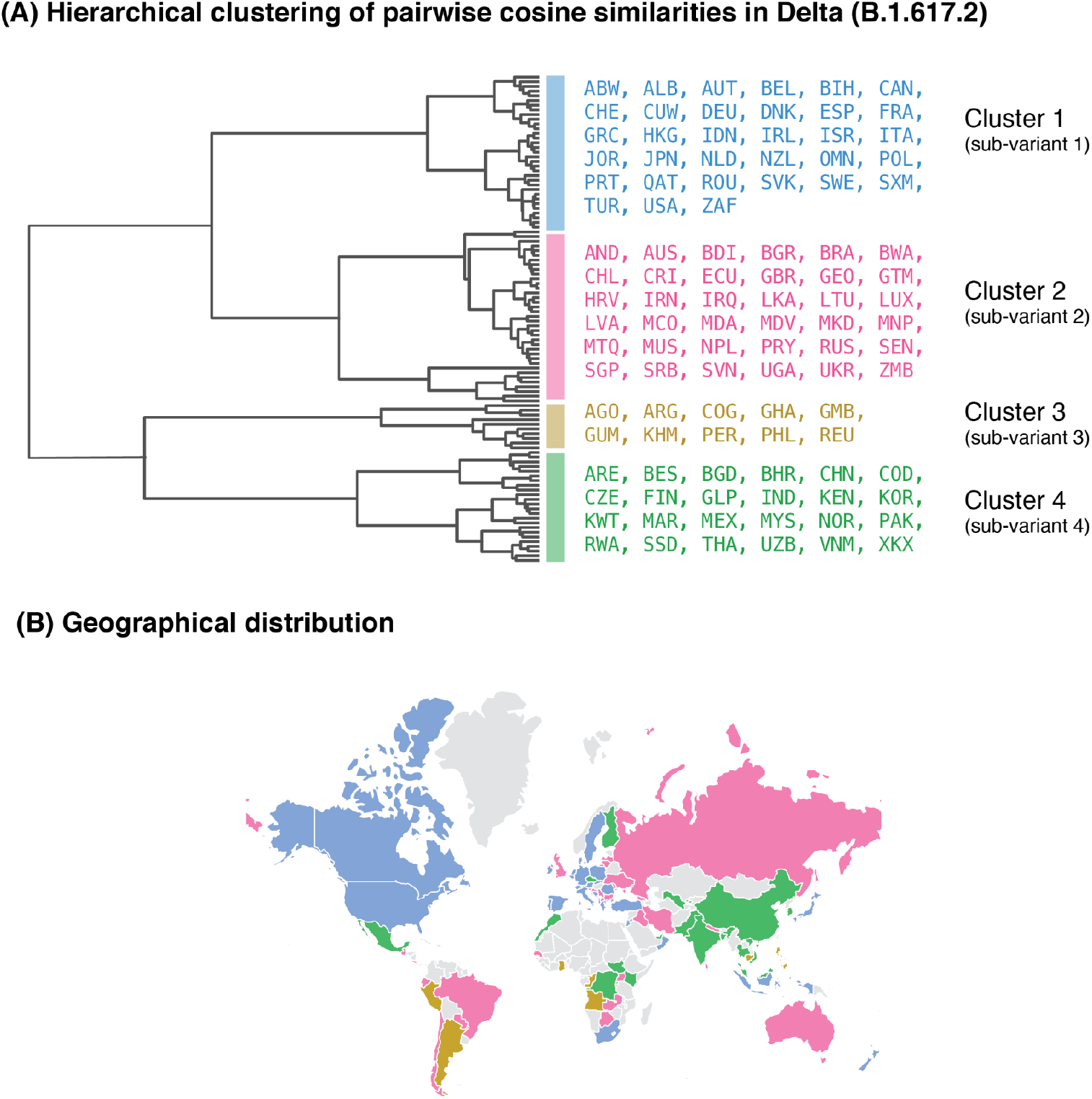
Comparison of the Delta sub-variants. **(A)** Hierarchical clustering of pairwise cosine similarities across countries. We identified four clusters corresponding to four sub-variants of the Delta variant. The dendrogram shows the hierarchical relationship among the Delta sub-variants. **(B)** Geographical locations of the countries of localization of the sub-variants. The annotations on a map of the world show that the sub-variants are prevalent in geographically distant countries.

**Figure S5:**
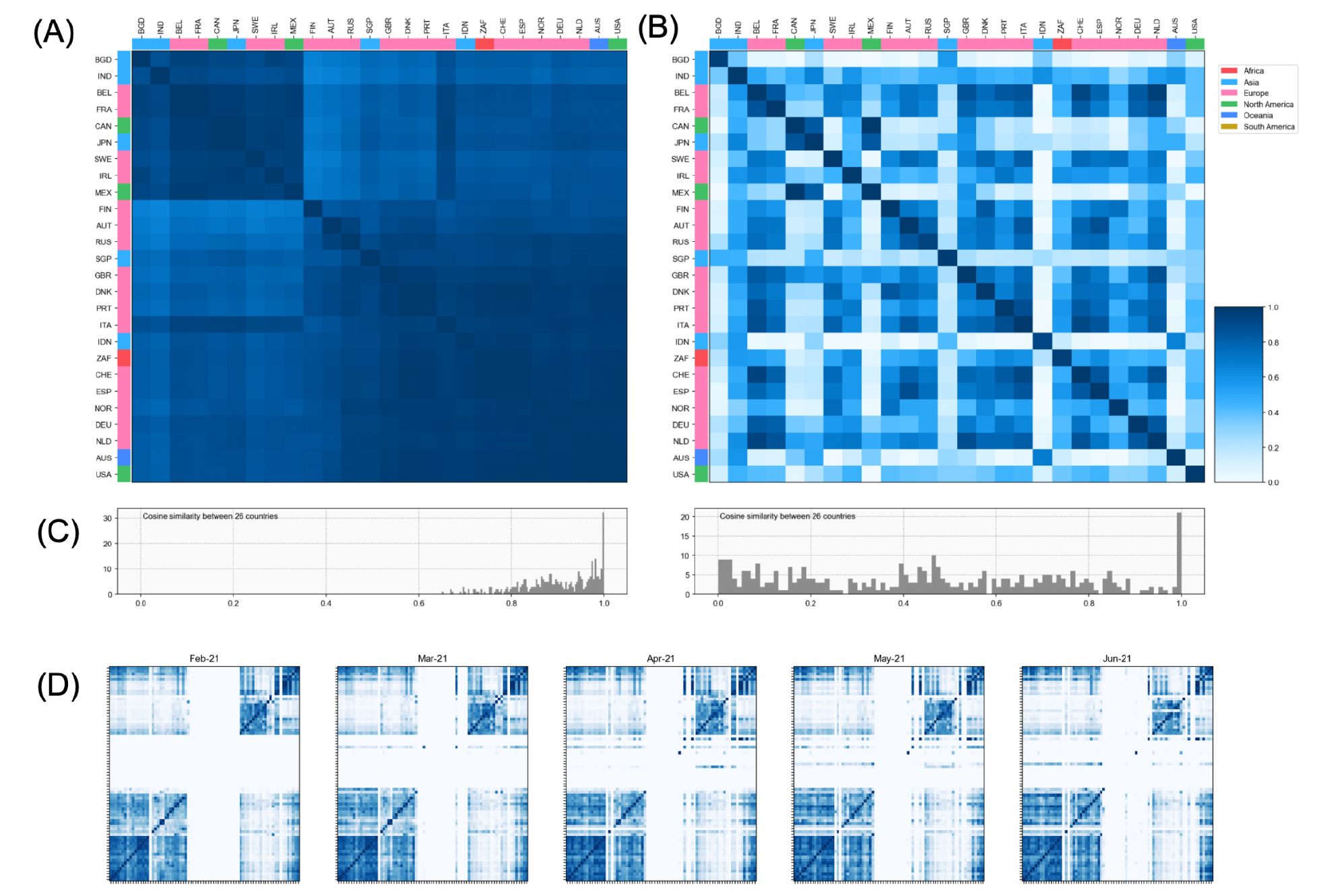
Effect of geographical separation and airline connectivity on the diversity of the Delta variant. **(A)** Cosine similarity between country-specific core mutations in Delta variant across all countries of its occurrence. This shows that the diversity of the Delta variant is not affected by the geographical separation of the countries. **(B)** Airline connectivity across countries. Sub-variants of the Delta variant may or may not co-exist in countries with good airline connectivity. This shows that the diversity of the Delta variant is not affected by airline connectivity across countries. **(C)** Frequency histogram for the distribution of the cosine similarity values. **(D)** Temporal airline connectivity. We observe that the patterns of airline connectivity have been unaffected during the period of February 2021 - July 2021.

**Figure S6:**
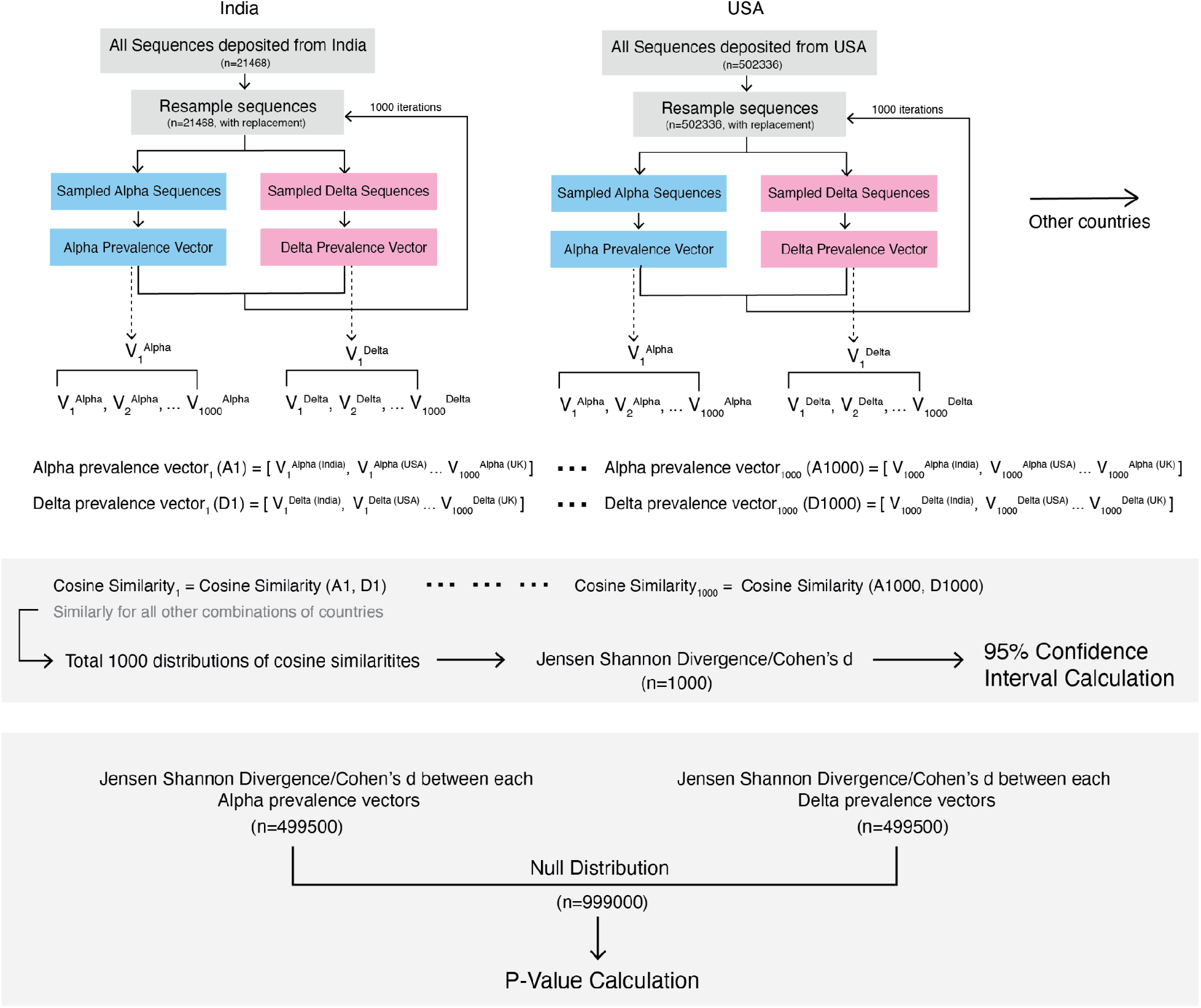
Bootstrapping Methodology. Methodological details of bootstrapping method.

**Figure S7:**
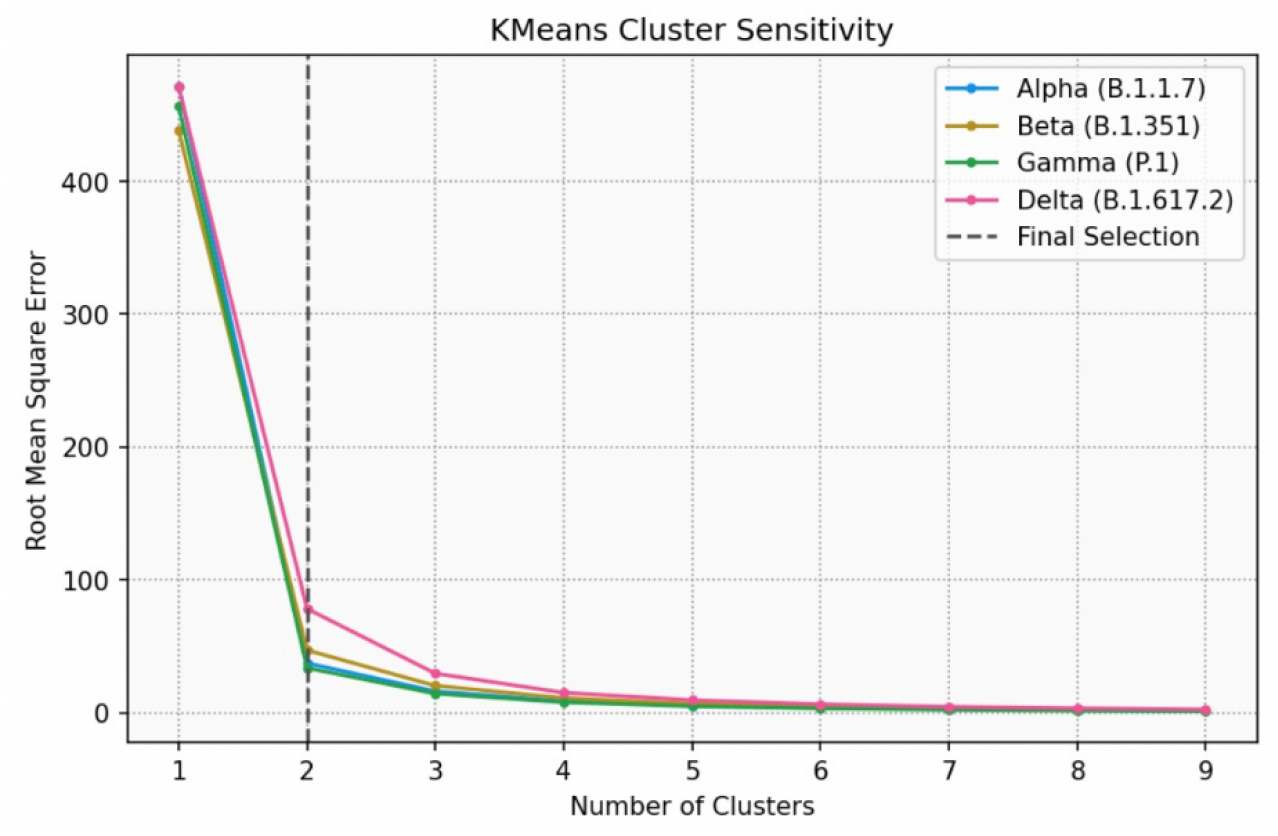
K-means Sensitivity analyses. K-means clustering sensitivity analysis. Country-specific core mutations for each variant of concern are calculated using a varying number of clusters (*k*) in the K-means algorithm. For each value of *k*, the Root Mean Square Error (RMSE) was calculated based on the distance of points from the centroid. The mean RMSE across all countries for each value of *k* has been plotted here. The dotted line represents the value of *k* (=2) selected for further analyses.

**Figure S8:**
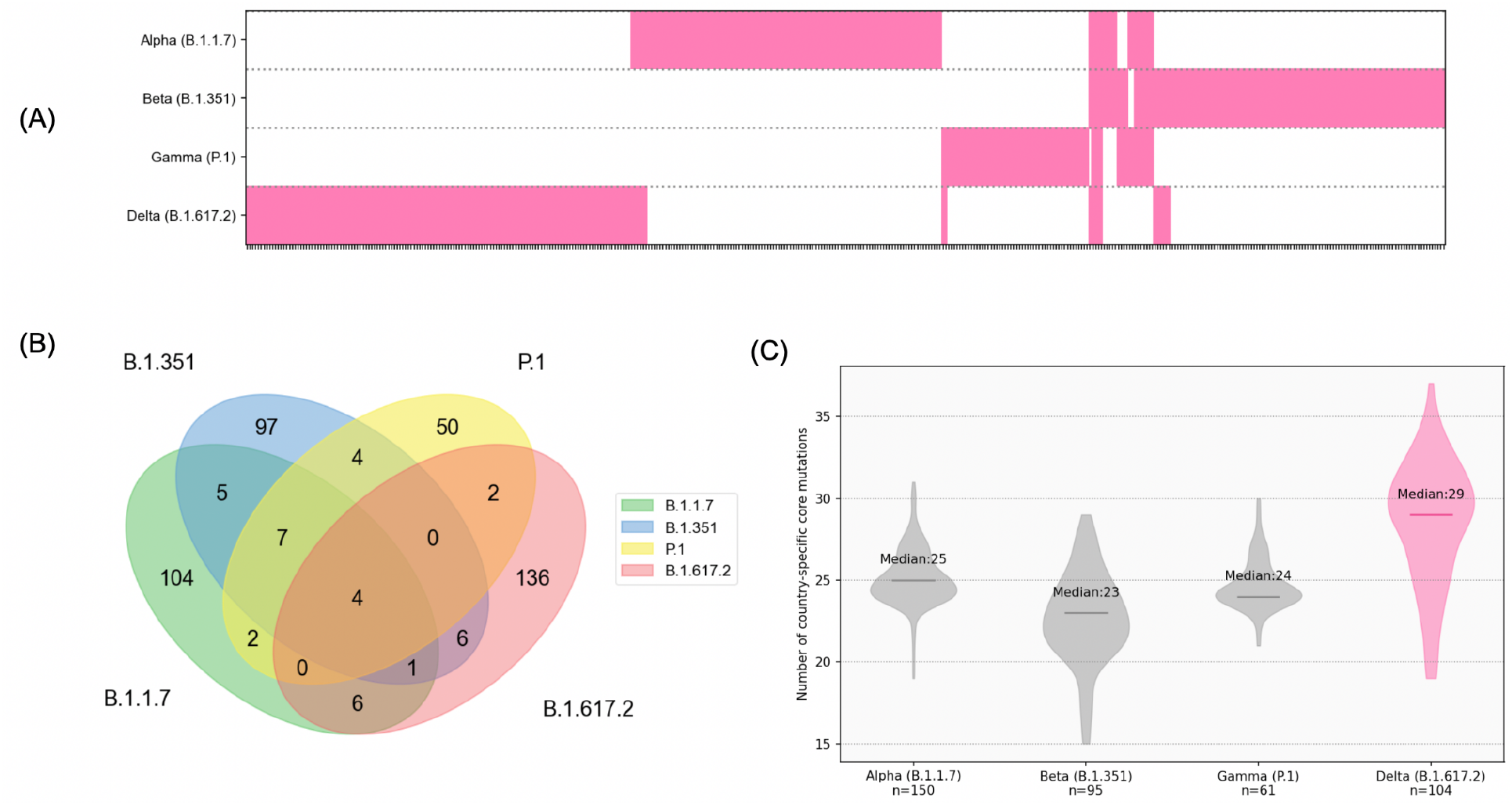
Distinct mutational profiles of the SARS-CoV-2 variants. **(A)** The different repertoire of mutations in the variants of concern. Each tick on the x-axis represents a mutation and is annotated in magenta if it is a part of the union set of country-specific core mutations for the variant. **(B)** Comparison of union sets of country-specific core mutations across different variants of concern. 136 out of 155 (87.74%) mutations in the Delta variant are unique to it. **(C)** Distribution of country-specific core mutation counts across countries for the variants of concern. The median count of country-specific core mutations for each variant is indicated on the plot. The Delta variant has a higher mutational load as compared to the other variants of concern.

**Figure S9:**
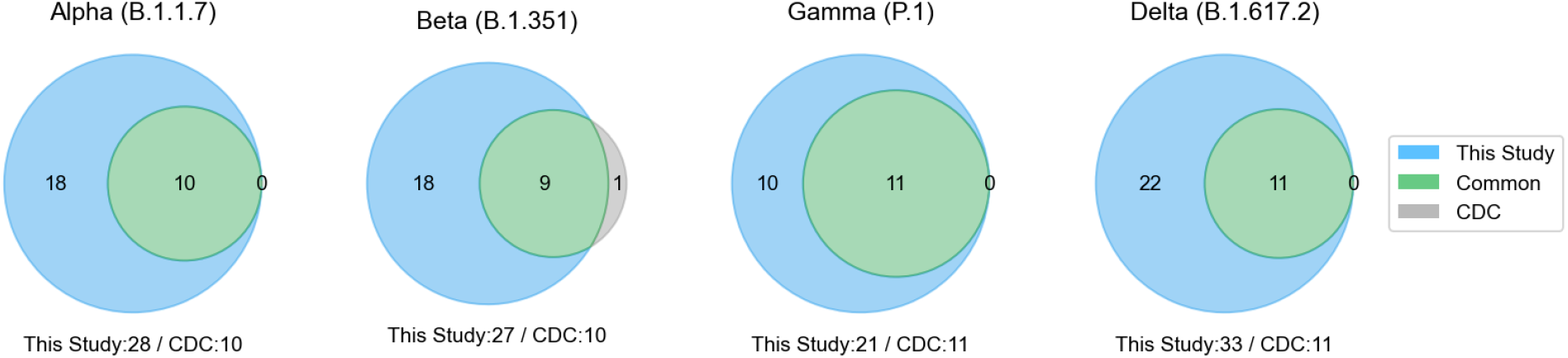
Union set of country-specific core mutations in Spike protein of SARS-CoV-2 variants. Comparison of country-specific core Spike mutations versus characteristic Spike mutations. Venn diagrams show a comparison of core Spike mutations identified in this study versus characteristic Spike mutations reported by CDC for each variant of concern. We failed to identify ΔL241 from the CDC ‘s set, which can be attributed to the differential alignment of the deleted positions to the wild-type SARS-CoV-2 genome in the mutation calling pipeline of the genomic assembly. In contrast, we identified multiple mutations in the Spike protein for each lineage that are highly prevalent in one or more countries but are missing from CDC ‘s set of mutations.

**Table S1:** Highly prevalent mutations in the Delta variant. The highly prevalent mutations in the Delta variant are listed here. Other than Spike D614G and NSP12 P323L, all the other mutations are nearly exclusive to the Delta variant.

|  | Prevalence (in %) |  |  |  |
| --- | --- | --- | --- | --- |
|  | Alpha (B.1.1.7)<br>(n=813,973) | Beta (B.1.351)<br>(n=177,46) | Gamma (P.1)<br>(n=42,869) | Delta (B.1.617.2)<br>(n=198,460) |
| M_I82T | 0.03 | 0.04 | 0.01 | 99.90 |
| NS3_S26L | 0.15 | 0.20 | 0.21 | 99.92 |
| NS7a_T120I | 0.15 | 0.08 | 0.09 | 99.72 |
| NS7a_V82A | 0.01 | 0.02 | 0.00 | 99.38 |
| N_D377Y | 0.05 | 0.20 | 1.00 | 99.63 |
| N_R203M | 0.00 | 0.02 | 0.00 | 99.87 |
| NSP12_P323L | 99.90 | 91.68 | 99.70 | 99.87 |
| Spike_D614G | 99.94 | 99.63 | 99.97 | 99.95 |
| Spike_L452R | 0.05 | 0.06 | 0.01 | 99.85 |
| Spike_P681R | 0.19 | 0.03 | 0.03 | 99.89 |
| Spike_T19R | 0.00 | 0.01 | 0.00 | 99.84 |
| Spike_T478K | 0.01 | 0.06 | 0.01 | 99.88 |

**Table S2:** List of the union set of country-specific core mutations unique to the Delta variant ‘s proteome. This table lists the union set of country-specific core mutations (136), across 104 countries, which are unique to the proteome of the Delta variant (as in **Figure S8B**).

| Protein | Mutations |
| --- | --- |
| Spike (S) | T19R, K77T, T95I, G142D, E156G, ΔE156, ΔF157, R158G, ΔR158, R214H, P251L, D253A, L452R, T478K, E484Q, T572I, D574Y, Q677H, P681R, D950B, D950N, D979E, G1167V, M1229I, D1259Y, V1264L |
| Nucleocapsid (N) | G18C, D63G, L139F, R203M, G215C, A252S, S327L, D377Y, R385K |
| Membrane (M) | D3H, I82T |
| NSP1 | E87D, ΔK141, ΔS142, ΔF143 |
| NSP2 | K81N, P129L, P200L, P200S, L204F, I251M, A596T |
| NSP3 | A85V, P192L, T422I, A488S, T678I, I707V, H727R, V765F, P778S, P822L, E906D, V1070I, P1103Q, T1189I, L1244F, G1273S, H1274Y, A1311V, P1469S, I1723V, A1736V, T1830I |
| NSP4 | S137L, V167L, T204I, D217N, V293I, C296F, F375S, T492I |
| NSP5 | I213V |
| NSP8 | K37N, L122I |
| NSP10 | A104V |
| NSP6 | A2V, H11Q, T77A, V149A, T181I |
| NSP12 | K91R, M124I, F192V, M196I, R197Q, G228S, D269G, D481A, T644M, G671S, T801I, M818V |
| NSP13 | P77L, V187L, V210I, I334V |
| NSP14 | T16I, P46L, M49I, M72I, A119V, V182I, I363V, A394V, Y420stop |
| NSP15 | K109N, H234Y, R257C, K259R, M330I |
| NSP16 | P215L, R216N, R216C, Q238H |
| NS3 | S92L, A110S, E239D |
| NS6 | K48N |
| NS8 | A65S, S69L, I88T, L95F |
| NS7a | P45L, V71I, V82A, V104F, L116F, T120I |
| NS7b | T40I |

